# Investigating the distribution of antibiotic resistance genes in relation to bacterial, fungal, and functional diversity in a hay field

**DOI:** 10.1101/2025.07.24.666709

**Authors:** Carolina Oliveira de Santana, Pieter Spealman, Conrad Vispo, David Gresham, Sage Saccomanno, Christopher N. LaFratta, Swapan S. Jain, Robert S. Dungan, Gabriel G. Perron

**Affiliations:** Department of Biological Sciences, State University of Feira de Santana, Feira de Santana, Bahia, Brazil; Bard Center for Environmental Sciences and Humanities, Bard College, Annandale-On-Hudson, NY, 12504; Center for Genomics and Systems Biology, New York University, New York, NY 10004, USA; Hawthorne Valley Farmscape Ecology Program, Hawthorne Valley Association, Ghent, NY, 12075, USA; Chemistry and Biochemistry Program. Bard College, Annandanle-On-Hudson, NY 12504; USDA-ARS, Northwest Irrigation & Soils Research Laboratory, Kimberly, Idaho 83341, USA

## Abstract

The spread of antibiotic resistance in pathogenic bacteria is one of the most pressing public health threats of the 21^st^ century, with at least 5 million deaths currently associated with it. While recent work has shown the importance of environmental reservoirs in the emergence of antibiotic resistance genes (ARGs), it is unclear which features of microbial diversity relate to ARGs of clinical relevance. Here, we investigate the role of bacterial, fungal, and functional diversity on the distribution of clinical and environmental ARGs along a single transect located in an aging hay field on an otherwise active farm. This transect spans a length of several hundred meters, increasing in distance from an agricultural access road and stream. We use *16S rRNA* and *ITS* amplicon sequencing to measure bacterial and fungal diversity respectively, in combination with whole-genome sequencing (WGS) to characterize functional and ARG diversity. We find increasing bacterial and functional diversity along the transect, as well as distinct community structures for both bacteria and fungi. While we find that the diversity of environmental ARGs is significantly correlated with both bacterial and fungal diversity, clinical ARG diversity only significantly decreased as fungal diversity increased. Our results suggest that while microbial diversity increases with distance from the road and stream; and this diversity tends to determine the diversity of the majority of ARGs; this trend is not observed for ARGs of clinical relevance, which appear to be largely driven by the variety of fungal groups in the environment.

## 1. Introduction

The spread of antibiotic resistance in pathogenic bacteria is one of the most pressing public health threats of the 21^st^ century (Davies et al. 2013; Levy and Marshall 2004). Current models state that roughly 5 million deaths can be associated with antibiotic-resistance bacteria globally and that this number could double by 2050 if no actions are taken (de Kraker, Stewardson, and Harbarth 2016; Antimicrobial Resistance Collaborators 2022). Even though stopping such infections in hospitals has become a major priority around the world (“Website,” n.d., *The Bacterial Challenge: Time to React : A Call to Narrow the Gap Between Multidrug-Resistant Bacteria in the EU and the Development of New Antibacterial Agents* 2009; Kahn 2016), recent work has revealed that novel antibiotic-resistant bacteria strains often emerge in environmental reservoirs (Aminov 2009; Finley et al. 2013; Kraemer, Ramachandran, and Perron 2019; Perron, Quessy, and Bell 2008). Environmental reservoirs, here, refers to those biotic and abiotic habitats that support microbial ecologies such that they are independent of the broader environment. Define environmental reservoirs? Indeed, antibiotic pollution resulting from medical, industrial, domestic and agricultural activities (Wilkinson et al. 2022; X. Li et al. 2018; Larsson 2014) combined with the release of antibiotic-resistant bacteria of public health concerns via wastewater and farming (Slekovec et al. 2012; Young, Juhl, and O’Mullan 2013; Fogler et al. 2019) create selective environments on an unprecedented global scale (Palumbi 2001; Smith, Feil, and Smith 2000; Perron, Inglis, et al. 2015). Yet, regulators in most countries pay little attention to the spread of antibiotic-resistant bacteria in the environment (Kahn 2016; Kraemer, Ramachandran, and Perron 2019).

Agricultural soil ecosystems are also increasingly recognized as an important reservoir of antibiotic-resistant bacteria (Fogler et al. 2019; Côté and Quessy 2005). In addition to hosting an impressive taxonomic diversity of bacterial and fungal microorganisms (Caporaso et al. 2011; Fierer and Jackson 2006), most soils harbor a vast diversity of antibiotic resistance genes, or ARGs, naturally present in a diversity of microorganisms (D’Costa et al. 2006; Allen et al. 2009). Indeed, due to the presence of antibiotic-producing microorganisms like *Streptomyces* spp., many antibiotic-resistance genes are present in soils; many of of these genes predate antibiotic use in modern medicine (D’Costa et al. 2011; Perron, Whyte, et al. 2015) and are unique to the soil environment (Forsberg et al. 2014; Gibson, Forsberg, and Dantas 2015; Forsberg et al. 2012). Yet, the diversity of antibiotic-resistance genes present in soil, combined with the ability of bacteria to acquire novel resistance genes horizontally, represent an important reservoir of resistance (Byrne-Bailey et al. 2011; Ghaly et al. 2019).

Indeed, the presence of clinically important antibiotic resistance, that is resistance genes that have been observed in clinical contexts, in pathogenic bacteria is higher in agricultural soil of all kinds compared to non-agricultural sites (Dungan, Strausbaugh, and Leytem 2019; Balta et al. 2025). Agricultural soils are frequently exposed to antibiotics and enteric bacteria, often carrying antibiotic resistance genes, via manure used as fertilizer, even if the manure is organic and has been composted (X. Li et al. 2018; Côté and Quessy 2005). Moverover, microbial communities in heavily farmed agricultural soils are exposed to extensive selective pressures exerted by intensive farming practices such as monoculture and the extensive use of pesticides and herbicides, which disrupt the stability of soil microbiomes (Marti et al. 2013; Demoling, Figueroa, and Bååth 2007; Cycoń, Mrozik, and Piotrowska-Seget 2019). Lastly, heavy metal concentrations have been identified as highly correlated with antibiotic resistance, independent of any specific land use such as agriculture (Pagaling et al. 2023). Under such conditions, ecological functions such as productivity and crop yield (Belete and Yadete 2023), organic matter decomposition and bioremediation are impacted (Prashar and Shah 2016) as well as a susceptibility to invasion (Chen et al. 2019). In other words, in addition to impacting ecosystem functions, the persistent antibiotics, pesticides, heavy metals, and other chemicals present in industrial agricultural settings give resistant invasive, opportunistic, or pathogenic soil bacteria an opportunity to flourish.

For this reason, agricultural soils represent an interesting opportunity to investigate whether farming practices that favor soil health and microbial diversity, such as crop rotation and no-till for example (Yoon et al. 2024), will also lead to a lower level of antibiotic resistance of clinical importance in the environment.

While it is possible that under certain conditions soil could show high levels of diversity due to the “Intermediate Disturbance Hypothesis” (Connell 1978; X. Zhang et al. 2018), healthy soils are generally characterized by a high diversity of bacterial and fungal microorganisms (Yoon et al. 2024). Of course, it is important to note that comparing soil microbial diversity across different ecosystems presents challenges due to varying environmental conditions, which complicate direct comparisons of microbial diversity and its implications for soil health across ecosystems (Kačergius et al. 2023).

Interestingly, due to their unique physiology, bacteria and fungi are also likely to modulate the presence of antibiotic resistance genes in idiosyncratic ways (Han et al. 2022). For example, fungi are more likely to secrete antimicrobials in the environment, which could have an important impact on the distribution of antibiotic resistance genes in soil where fungi are abundant (Hashmi et al. 2017). Inversely, because many bacteria harbor resistance to antibiotics naturally occurring in the environment, we might expect high bacterial diversity to correlate with high diversity in genes resistant to those naturally occurring antibiotics. Similarly, soil under heavy human influence may feature a higher relative abundance of antibiotics from anthropogenic sources and therefore a higher proportion of clinically-relevant ARGs (Forsberg et al. 2014; Osburn et al. 2023). In fact, functional redundancy often observed in diverse ecosystems, where multiple species perform similar roles, can stabilize ARG prevalence, even under disturbance (Allison and Martiny, n.d.; Avila-Jimenez et al. 2020).

Here, we test the hypothesis that antibiotic-resistance genes (ARGs) of clinical importance will be less abundant in soil harboring more diverse microbial communities. Specifically, we test the prediction that antibiotic-resistance genes of clinical importance will be negatively correlated with bacterial diversity or with fungal diversity or both within a given ecosystem. Using *16S rRNA* and *ITS* amplicon sequencing to measure bacterial and fungal diversity respectively, in combination with whole-genome sequencing to characterize the abundance of antibiotic resistance genes, we investigate the distribution of antibiotic resistance genes in relation to bacterial, fungal, and functional diversity along a single transect located in a hay field left untouched for 5 years on an otherwise active farm.

## 2 Methods and Materials

### 2.1 Soil collection

Samples were collected from four sites along a transect perpendicular to Esopus Creek, NY, as described in de Santana et al. (2023) from Field 8 of Hudson Valley Farm Hub. The latter is a hay field seeded in 2017 with a custom seed mix composed of Perennial Rye, Annual Rye, Timothy, Orchard Grass, and Huia Clover (a White Clover). Prior to this, the field was predominantly planted with oats and clover in 2016; supported vegetables in 2015, and earlier records suggest a history of sweet corn cultivation pre-dating its integration into the Farm Hub system. Management practices for the hay field have included annual fall mowing, with the exception of 2023, when mowing was deferred to address habitat considerations for avian species.

The four sites, chosen along a transect in increasing distance from a road and waterway, were labeled as follows: F8-W, a forested strip between the riverbank and a dirt road used by farming equipment; F8-0 (3.05 m), located in the hay field immediately next to the dirt road and about 15 feet from F8-W’s forest edge. F8-300, located 300 feet (91.44 m) west of the forest edge and F8-600, located 600 feet (182.88 m) into the hay field. At each site, we collected three samples of 10 g topsoil (∼5 cm deep) each, from sample sites located within 1 m of each other running parallel to the stream using sterile techniques. We sampled the transect three times over the spring and summer: 6 June, 19 June, and 3 July 2019. Samples were transposed to the laboratory on ice in a dark cooler and frozen for at least 24 h before being processed. We geolocated one point along the transect at each distance to ensure that the exact locations were being sampled each time using GPS coordinates.

### 2.2 DNA extractions and sequencing

We extracted and purified DNA from 1.8 g of soil for each sample using the Quick-DNA Fecal/Soil Microbe Miniprep kit (Zymo Research, Irvine, CA, USA). We then used this DNA for 16S rRNA and internal transcribed spacer (ITS) amplicon sequencing as well as whole-genome the V4 regions using the 515F and 806R primers as per the Earth Microbiome Project (Walters et al. 2016). For ITS amplicon sequencing, we used the ITS1f and ITS2 primers (Bokulich and Mills 2013). PCR products for 16S rRNA and ITS amplicons were pooled separately and purified on a 2% agarose gel using the Qiagen gel extraction kit. As for whole-genome sequencing (WGS), libraries were prepared using the Nextera XT DNA Library Preparation kit (Illumina) and gel-purified. All purified libraries were quality checked using an Agilent 2100 Bioanalyzer and DNA High Sensitivity kit and pooled in an equimolar ratio. Purified pools were then stored at −20 °C until sequencing. Amplicon libraries were sequenced at Wright Labs, Huntington, PA) using Illumina MiSeq v2 paired-end sequencing (2 × 250-bp reads) with 20% PhiX spike-in. WGS libraries were also sequenced at Wright Labs using an Illumina NextSeq 2000 (2 × 150-bp paired-end reads) with default parameters.

### 2.4 Amplicon reads pre-processing and analysis

For *16S rRNA* and *ITS* amplicons, demultiplexed sequence reads were filtered and trimmed with Trimmomatic (Bolger, Lohse, and Usadel 2014) (ILLUMINACLIP:TruSeq3-PE.fa:2:30:10 LEADING:3 TRAILING:3 SLIDINGWINDOW:4:15 MINLEN:100) with the requirement of a minimum average read quality score of 15 for inclusion. For each read, the sliding window cuts any read at the point where the median quality score over a 4nt window is less than 15. To identify amplicon sequence variants or ASVs, that is unique identifiable sequences, we used the QIIME2 pipeline (v2021.2) with default parameters except for DADA2 (16S rRNA, denoise-paired, –p-trim-left-f 0 –p-trim-left-r 0 –p-trunc-len-f 250 –p-trunc-len-r 250; ITS, denoised-single, –p-trunc-len 150). All ASVs were retained in the dataset. Filtering was performed only on taxa and only for differential abundance analysis. Taxonomic assignment was performed using QIIME2’s naive Bayes scikit-learn classifier (Bokulich et al. 2018) trained using the 16S rRNA gene sequences in the SILVA database (Silva SSU 138)(McDonald et al. 2012) and UNITE’s dynamic all taxa database v8.3 for ITS (Abarenkov et al. 2021). Two ITS samples, F8-300 (2019-06-19) and F8-600 (2019-07-03), had too few reads and were removed from further analysis (Supplemental Table S1).

Using ASVs as the unit of taxonomic identification, we estimated the total number of observed taxa (Supplemental Table S2). For the purposes of alpha- and beta-diversity tests we used QIIME2 rarefied 16S (max = 10K) and ITS (max = 350K) abundances. For all alpha-diversity comparisons between sites significance was evaluated using Kruskal-Wallis (KW) statistics with adjusted *P*-values (Supplemental Table S3). For all beta-diversity comparisons between sites significance was evaluated using PERMANOVA statistics (Supplemental Table S4).

We characterized the variance between time points and found no overall statistical effect of sampling date on bacterial diversity, measured either as the total number of observed bacterial ASV’s (*H*_(2)_ = 2.14; *P* = 0.34) or Shannon’s diversity index (*H*_(2)_ = 0.63; *P* = 0.73). Furthermore, we did not observe statistical differences between bacterial population structure at different time points using Bray-Curtis dissimilarity index (adonis: pseudo-*F*_(2,30)_ = 1.0743; *R*^2^ = 0.06; *P* = 0.32). We also did not observe significant differences between dates in the total number of observed fungal ASV’s (*H*_(2)_ = 0.3; *P* = 0.98) or Shannon’s diversity index (*H*_(2)_ = 2.13; *P* = 0.34). Fungal community structure between sampling dates, based on Bray-Curtis dissimilarities, also did not change significantly (adonis: pseudo-*F*_(2,30)_ = 0.7167; *R²* = 0.05; *P* = 0.88). With this degree of similarity in mind we treat these time points as biological replicates hereafter.

Furthermore, we used the Phyloseq package (McMurdie and Holmes 2013) for advanced analysis of sample subsets and specific taxonomy levels, and ggplot2 (Wickham, 2009) for the visualization of data. We used the Vegan package (Dixon 2003) for the diversity analysis of prokaryotic and fungal communities as well as ARGs and functions. Vegan was also used for permutation analysis to identify differences between taxonomic and functional structures, as well as the resistome at each site. We used Kruskal-Wallis test in R (version 4.2.1) (“The R Project for Statistical Computing,” n.d.) to check for statistical differences in the abundance of ARGs between collection sites and as an effect of the water influence. Correlation analysis was performed using in-built functions in the R environment using linear model (“lm”) on the data of diversity.

### 2.5 Whole-genome sequencing read processing and analysis

In order to access the taxonomic and functional profiles of the prokaryotic communities in the whole-genome sequencing, we used the MGnify 5.0 pipeline (Mitchell et al., 2019a) from the European Nucleotide Archive (ENA) online platform (https://www.ebi.ac.uk/ena). Reads were assembled with metaSPAdes (v3.15.3) (Nurk et al., 2017). Prediction of genes was performed using Prodigal (v2.6.3) (Hyatt et al., 2010) and FragGeneScan (1.20) (Rho et al., 2010), while functional annotation was performed with InterPro (v75.0) (Mitchell et al., 2019b), a custom implementation of Gene Ontology (Ashburner et al., 2000), and Kegg Ortholog (v90.0) (Kanehisa et al., 2016) using Kofam (v2019-04-06) (Aramaki et al., 2015). Individual proteins were then compared against UniRef90 (v2019_11) (Suzek et al., 2015) using DIAMOND (v0.9.25.126) (Buchfink et al., 2015).

Furthermore, the WGS sequences were assembled using MEGAHIT (D. Li et al. 2015) on the Galaxy platform (Jalili et al. 2020) and the resulting contigs were then downloaded and locally compared to the CARD database (Alcock et al. 2019) using the RGI tool in order to calculate the ARGs present within these contigs. In addition, we used (Z. Zhang et al. 2022) to classification of health risk ARGs to assign high-risk, clinically-relevant ARGs (eg. Q1) or low risk ARGs (Q2+). The latter is indicative of resistance genes common in the environment (here, referred to as “environmental ARGs”) and unlikely to be linked to clinical issues while the former indicate resistance genes currently linked with clinical implications for health (here, referred to as “clinical ARGs”).

To visualize the correlation between ARGs and predicted gene function we first subset both groups to only include instances that present in a minimum of 4 samples (build_linregression.py). We next ran a linear regression on all pairwise combinations of normalized abundances, generated by RGI for the ARG or InterPro for gene function, the resultant p-values were then corrected using Benjamani-Hochberg (BH). The heatmap shows those pairwise combinations with BH adj.p-values <= 0.05.

### 2.6 Quantifying the abundance of specific antibiotic resistance genes

The DNA extracts were analyzed by quantitative real-time PCR (qPCR) on a CFX96 Touch Real-Time PCR Detection System (Bio-Rad, Hercules, CA) according to methods described in full elsewhere (Dungan, McKinney, and Leytem 2018). In brief, each individual reaction was performed with SsoAdvanced™ Universal SYBR Green Supermix (Bio-Rad). In the present study, the gene targets were 1*6S rRNA*, *aadA*, *ampC*, *bla*_CTX-M_, *erm*(B), *intI1*, strA, *sul1*, *tet*(M), and *vanA*. The qPCR runs included a standard curve covering seven orders of magnitude, and each sample was analyzed in duplicate. Standards were created using gBlocks^TM^ Gene Fragments (Integrated DNA Technologies, Coralville, Iowa, USA). Primers and annealing temperatures for *blaCTX-M, erm(B), intI1, sul1,* and *tet(M)* are described in Dungan et al. (2018); conditions for genes *aadA*, *ampC*, *strA* and *vanA* are described in Supplementary Table S5. After estimating the copy number for each gene using the standard curves, we calculated the relative abundance of each gene by dividing their copy number by the copy number of *16S rRNA*, the latter being a bacteria universal gene used as a proxy for total bacteria count. For each gene present at all sampling sites, we performed statistical analysis in *R* by comparing models resulting from analysis of variance and linear regression, using the gene relative abundance as the dependent variable and sites as the independent variable.

### 2.7 Measuring heavy metal concentrations in soil samples

To evaluate the potential impact of pollution on Field 8, we measured the concentration (mg/kg) of eight heavy metals, i.e., Arsenic (As), Barium (Ba), Cadmium (Cd), Chromium (Cr), Copper (Cu), Nickel (Ni), Lead (Pb), and Zinc (Zn) at three replicates at all four sites and for two depths, i.e., 5 cm and 15 cm. First, ∼10 g of each sample was sent to the Cornell Soil Health Laboratories (Ithaca, NY) for standardized analyses (Moebius-Clune et al., n.d.). Then, we used another ∼5 g from the same collected soil to measure the concentrations of Mercury (Hg) using an in-house procedure. In brief, dried soil samples were ground with a mortar and pestle, then sieved using a USS #10 sieve. Approximately 1.0 g of soil (dry weight) was added to a 250 mL round bottom flask and refluxed for 15 minutes with 2.5 mL conc. HNO_3_ and 10 mL conc. HCl. After filtration through Whatman No. 41 filter paper, the filtrates were collected in 100 mL volumetric flasks. The filter was washed with 5 mL hot (∼95°C) conc. HCl and 20 mL hot reagent water, with the washings added to the same flask. Residues and filters were returned to the flask, refluxed with 5 mL conc. HCl, and reheated at 95°C until the filter dissolved. The solution was filtered again, cooled, and diluted to volume. The samples were centrifuged at 4600 rpm for 5 minutes, and the supernatants were transferred to scintillation vials. Heavy metal concentrations (mg/kg) were determined using the Agilent 5100 ICP-OES system (Agilent Technologies, Santa Clara, CA) after creating a standard curve. We then compared the concentrations of metals between the sites using analysis of variance, using the Tukey test for detecting differences between sites, when including the F8-W site. After removal of the F8-W site as an outlier, linear regression was used for testing the gradient of concentrations in the hay sites. We used a nested-anova to confirm that there was no difference in metal concentrations between samples taken at the surface, i.e., 5 cm, and soil samples at 15 cm.

## 3. RESULTS

### 3.1 Preliminary analysis of an unused hay field

To investigate the distribution of antibiotic resistance genes across the transect, we first sought to characterize the distribution of nine common antibiotic resistance genes relative to the abundance of the *16s rRNA* gene using qPCR. From the three genes that were present in all investigated sites, we found that both *aadA* (*F*_(1,7)_ = 20.16; *P* = 0.003; Supplemental Figure S1A) and *bla*_CTX_ (*F*_(1,7)_ = 8.39; *P* = 0.02; Supplemental Figure S1B) actually decreased in relative abundance with distance; while *vanA* did not show any correlation with distance (*F*_(1,7)_ = 0.09; *P* = 0.77; Supplemental Figure S1C). This trend, combined with the fact that overall microbial diversity is considered a sign of soil health within an ecosystem, suggests that there might exist a gradient of environmental pollution within the transect, with higher environmental pollution predicted to be higher near the field’s edge where it can be affected by moving vehicles and/or the creek.

To address this, we looked for the presence of nine heavy metals often associated with pollution. Heavy metals do not degrade over time and unlike other contaminants, they accumulate in soil and in living organisms. After analyzing the soil content for heavy metal concentrations we observed that, indeed, three out of nine metals were slightly but significantly higher near the edge of the field and decreased linearly with distance from the edge: Arsenic (*F*_(1,10)_ = 28.70; adj-*P* < 0.001; Figure S2A), copper (*F*_(1,10)_ = 87.27; adj-*P* < 0.0001; Figure S2C), and lead (*F*_(1,10)_ = 26.23; adj-*P* < 0.001; Figure S2D). In all other cases, the heavy metals did not show any significant change with distance. Interestingly, we also found that the metal concentrations at the site near the water, i.e., F8-W, was similar to that of the site near the field’s edge (Tukey test; *P*s < 0.5), except for copper which was lower near the water and lead which was higher (Tukey test; *P* < 0.5). Taken together, these results strongly suggest that there exists an environmental gradient within the transect, likely correlated with traces of environmental pollution in the soil.

### 3.2 Microbial diversity and community structure differ among sites along the transect

We then investigated the microbial diversity and community structures among the transect sites. We found that the average bacterial diversity significantly differed between sites (Pielou’s evenness: omnibus, KW, *P* = 0.014; Figure 1A). The highest average Pielou’s diversity index, accounting for taxa evenness in addition to the number of taxa, was found at site F8-600, located 600 ft into the hay field, indicating that the latter showed a more even distribution among bacterial ASVs than any other sites (adj-*P*s < 0.05).

**Figure 1.**
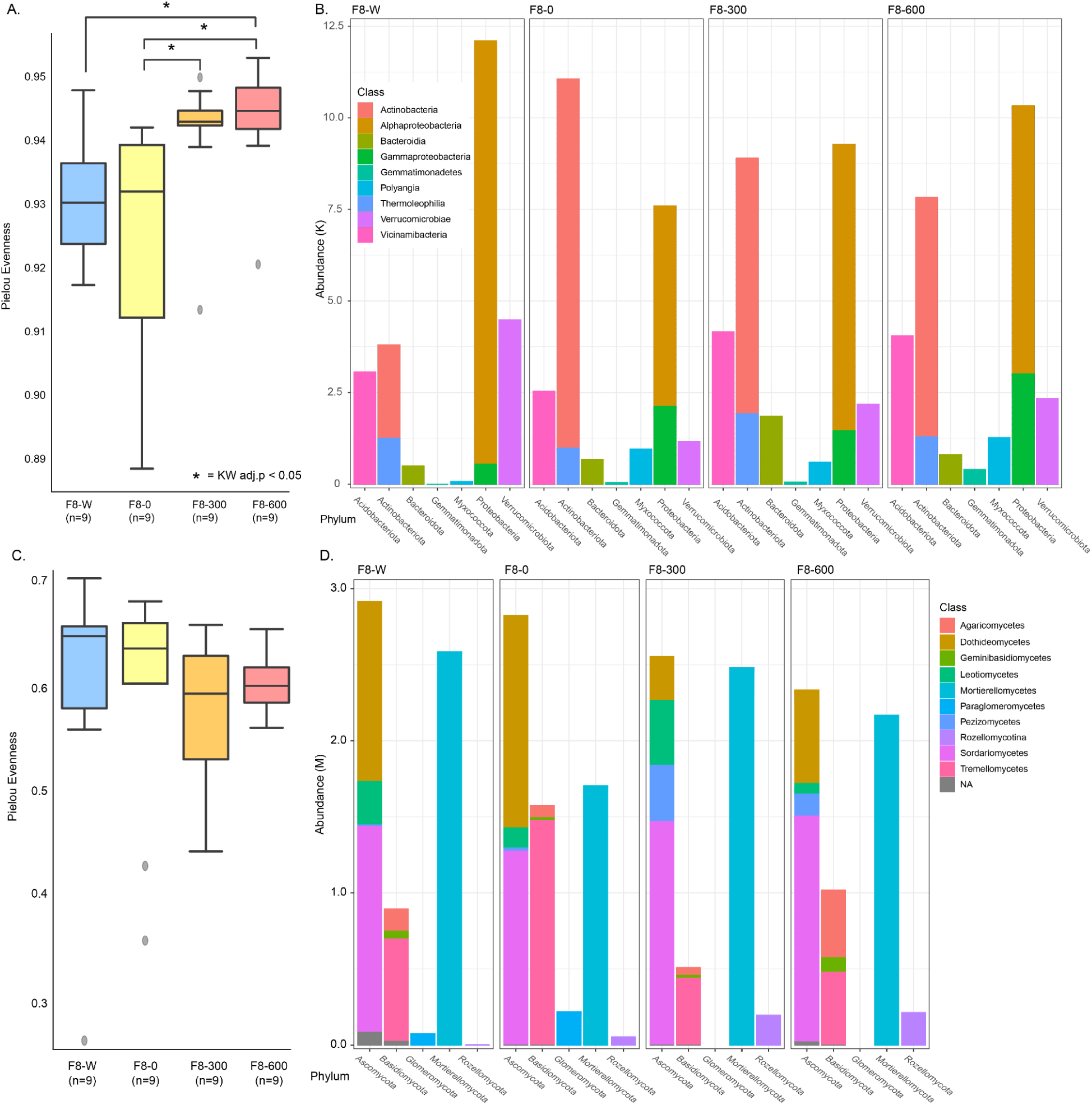
Description of microbial communities along the F8 agricultural transect. 16S rRNA diversity in each sample as measured by **A**) Peilou’s evenness, significantly different pairwise sites (KW, BH adj.P < 0.05) are indicated by an asterisk. **B**) Taxonomic box plot of the nine highest abundance 16S Classes, separated by phylum. ITS diversity in each sample as measured by **C**) Peilou’s evenness (no significant difference) **D**) Taxonomic box plot of the 10 highest abundance fungal Classes, separated by phylum.

We also found that bacteria community structures (Figure 1B) differed significantly between sites (Bray-Curtis: omnibus, PERMANOVA; *P* = 0.001; Supplemental ST4) as well as each site in a pairwise comparison (pairwise, PERMANOVA: adj-*P*s < 0.01), suggesting that these bacterial communities are distinct. Notably, the F8-W site had a median 20.5% similarity (similarity = 1-Bray-Curtis index) with the other three sites (Supplemental ST4) compared to a median of 36.3% within the other three sites, potentially because of the ecosystem differences between the forested F8-W site and the three hay field sites. In fact, when looking at community structure, communities found at each site along the transect were more closely related to each other, with an average 42.1% similarity, than to any other communities at other sites, with an average 27.8% similarity between the sites, suggesting that microbial communities might be impacted by an environmental gradient. Following taxonomic assignment, we found that the phyla Proteobacteria (median 33%) and Actinobacteria (median 20%) were the most abundant groups in all the collection sites.

Unlike what we found with bacterial communities, we did not find significant changes in Pielou’s evenness index for fungal ASVs (omnibus, KW, *P* = 0.433; Figure 1C). When comparing fungal community structures, however, we found significant differences between the four sites (Bray-Curtis: omnibus, PERMANOVA, *P* = 0.001; Supplemental ST4). Using Bray-Curtis dissimilarity index, we found that sites differed significantly from each other by an average of 62%, mainly due to significant differences between the sites F8-W and F8-300, suggesting again that F8-W is more distinct. We found that phyla Ascomycota (median 41%) and Mortierellomycota (median 26%) were the most abundant groups in all sites (Figure 1D).

As the preliminary analysis showed that the site F8-W was the most distinct for both the fungal and bacterial communities, we consider that the forested state and the close proximity to the water and the potential contamination present therein of herbicide, pesticide, and fertilizer runoff, in addition to it being forested make it a significant outlier, and therefore, the F8-W site is held out from the remainder of this analysis.

In addition to changes in diversity indexes in relation to distance we also sought to identify genera whose abundances correlated strongly with distance (Supplemental Figure S3). While we did identify numerous genera with statistically significant changes in abundance in a linear relationship to distance the vast majority of these had very small effect sizes (eg. change in relative abundance < 1% global abundance).

### 3.3 Functional diversity differs among sites along the transect

We also investigated total genetic diversity patterns using genes predicted from whole genome sequencing (WGS) data. Overall, we found 6,924 functions belonging to 207 metabolic pathways. We found the most common functions to be associated with transposase activity, along with common metabolic processes such as carbohydrate metabolic process and response to stress. We also found a number of metabolic pathways that are strongly associated with soil microbial communities such as regulation of nitrogen use, which has a great impact on soil fertility and ecosystem functioning, as the plants are incapable of nitrogen fixing.

An analysis of functional diversity showed that the number of observed genes differed between the three sites (Figure 2A). The genetic composition also showed significant differences between the sites, considering all sites (omnibus, PERMANOVA test: *P* = 0.035). When looking at functional diversity as a linear function of distance, we found that the number of different observed functions increased with distance (*F*_(1,7)_ = 7.62; *R*^2^ = 0.45; *P* = 0.02; Figure 2A). Similarly, we found that functional diversity, measured as the Shannon’s diversity index, increased significantly with distance away from the field’s edge (*F*_(1,7)_ = 10.01; *R*^2^ = 0.53; *P* = 0.02; Figure 2B). We also considered if functional diversity was correlated with either bacterial or fungal diversity; and while we did not find any statistically significant correlations between bacterial and functional diversity, we did find a significant negative correlation functional diversity and fungal diversity (Supplemental Figure S4) suggesting that fungal diversity may play a role in functional diversity.

**Figure 2.**
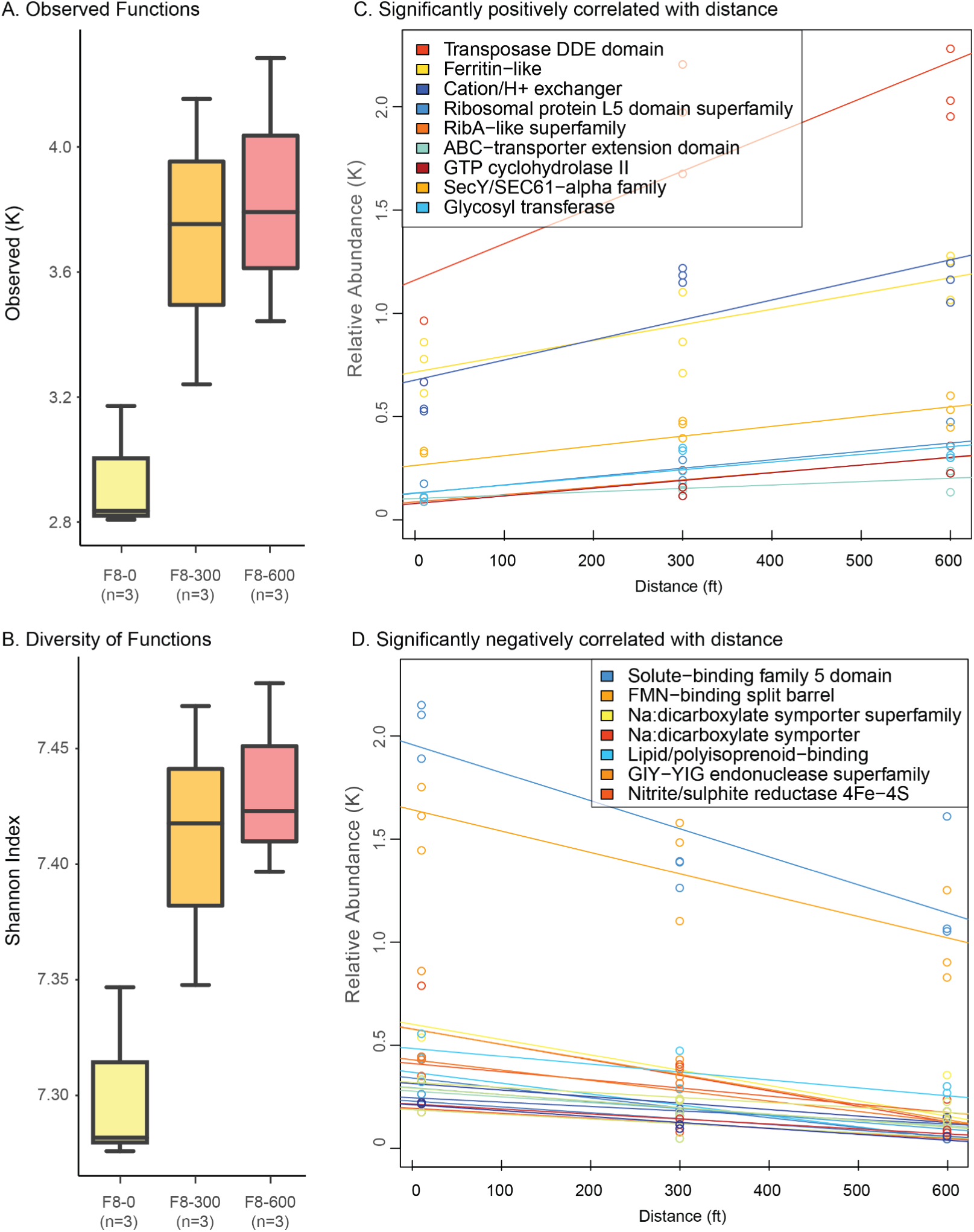
Description of genetic diversity along the F8 agricultural transect. Genetic diversity in each sample, i.e., individual black circles is represented as **A**) the number of Observed predicted genes, and **B**) Shannon diversity index in the soil samples per distance in the transect. Relationships between predicted individual functions and distance along the transect in the hay field were analyzed using linear regression and found to be **C**) positive (P < 0.0001), and **D**) negative (Ps < 0.0001) where individual circles represent data points for each gene at each site.

Lastly, we also tested for linear trends in all functions individually. Overall, 16 functions were significantly correlated with distance (Wald Test, BH adj.P <= 0.05). Some functions increase in abundance with distance (Figure 2C), such as the ABC-transporters; a family of exobiotic efflux transporters, well known for their role in providing multi-drug resistance in bacteria (Orelle, Mathieu, and Jault 2019) and potentially indicating a larger abundance of antibiotics in the soil and an increase in soil health. Conversely, numerous functions with distance; such as GIY-YIG endonuclease superfamily which encodes enzymes that repair double strand breaks and maintain genome stability under stress (Dunin-Horkawicz, Feder, and Bujnicki 2006), which could also indicate increased soil health.

### 3.4 Distribution of antibiotic resistance genes among sites along the transect

We next sought to investigate the distribution of antibiotic resistance genes across the transect. To do so, we further investigated the whole-genome sequencing data to look specifically for predicted antibiotic resistance genes (ARGs), with specific interest in high-risk, clinically-relevant ARGs (clinical ARGs) that have been previously identified (Z. Zhang et al. 2022) as compared to lower risk, non-clinical ARGs (environmental ARGs). Overall, we found 231 classes of ARGs, including 21 genes identified as high-clinical risk belonging to 5 classes. We found that the number of environmental ARG only changed marginally between sites (*H*_(2)_ = 5.4; *P* = 0.06; Figure 3A, blue) and the same was observed when considering ARG diversity estimated with Shannon index (*H*_(2)_ = 5.6; *P* = 0.06; Figure 3B, blue). Similarly, we found that the number of clinical ARGs did not change between sites (*H*_(2)_ = 1.28; *P* = 0.53; Figure 3A, red) nor showed different levels of diversity (*H*_(df)_ = 1.15; *P* = 0.56; Figure 3B, red).

**Figure 3.**
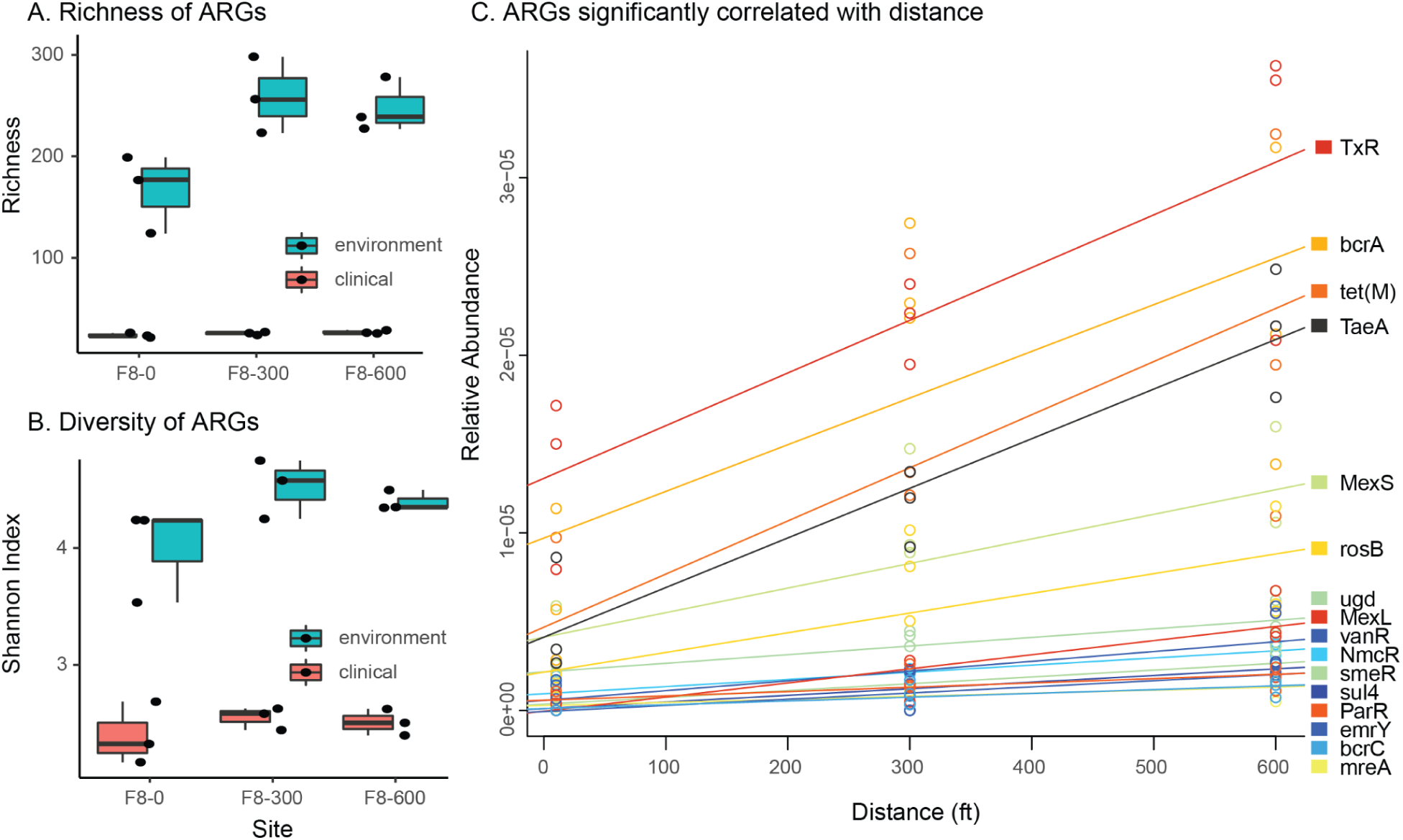
Description of antibiotic resistance diversity along the F8 agricultural transect. Total antibiotic resistance genes (blue) and clinical ARGs (red) diversity in each sample, i.e., individual black circles is represented as **A**) the number of Observed predicted ARGs genes, and **B**) Shannon diversity index in the soil samples per distance in the transect. Blue shade shows values in the wooded samples nearer the water. Data for each sampling date are presented separately but were analyzed as biological replicates. Relationships between predicted individual functions and distance along the transect in the hay field were analyzed using linear regression and found to be **C**) positive (P < 0.0001) where individual circles represent data points for each ARG at each sample site.

We then tested whether individual ARGs correlated with distance from the field’s edge by fitting a linear trend in the three hay field sites. We found 16 ARGs, detected in the majority of samples, to be significantly correlated with the transect distances (Wald Test, adj-*P* <= 0.05; Figure 3C). All of these significant ARGs that correlated with the distances in the transect increased relative to the distance from the road. Only three of these genes, tetM, ugd, and emrY are clinical ARGs, 1.2 fold lower number of observations than what would be expected by chance but not significantly so (FET, P=0.83).

### 3.5 Investigating relationships between antibiotic resistance and microbial diversity

One of the main objectives of this study was to investigate the distribution of antibiotic resistance genes across the transect in relation to bacterial and fungal diversity, especially when it comes to clinically relevant resistance genes. Based on previous research, we predicted that more diverse soil communities would correlate with a reduction in clinically-relevant ARGs.

We first investigated whether bacterial communities’ diversity could explain, at least in part, the distribution of antibiotic resistance in our samples. We found that neither the environmental ARGs abundance (*F*_(1,6)_ = 2.00; *R*^2^ = 0.44; *P* = 0.21; Figure 4A) nor the clinical ARG abundance correlated with the total number of bacterial ASVs (*F*_(1,6)_ = 0.24; *R*^2^ = 0.14; *P* = 0.64; Figure 4A). Similarly, we found that neither the environmental ARGs (*F*_(1,6)_ = 0.79; *P* = 0.79; Figure 4B), nor the clinical ARGs correlated with the total number of fungal ASVs (*F*_(1,6)_ = 3.53; *P* = 0.11; Figure 4B).

**Figure 4.**
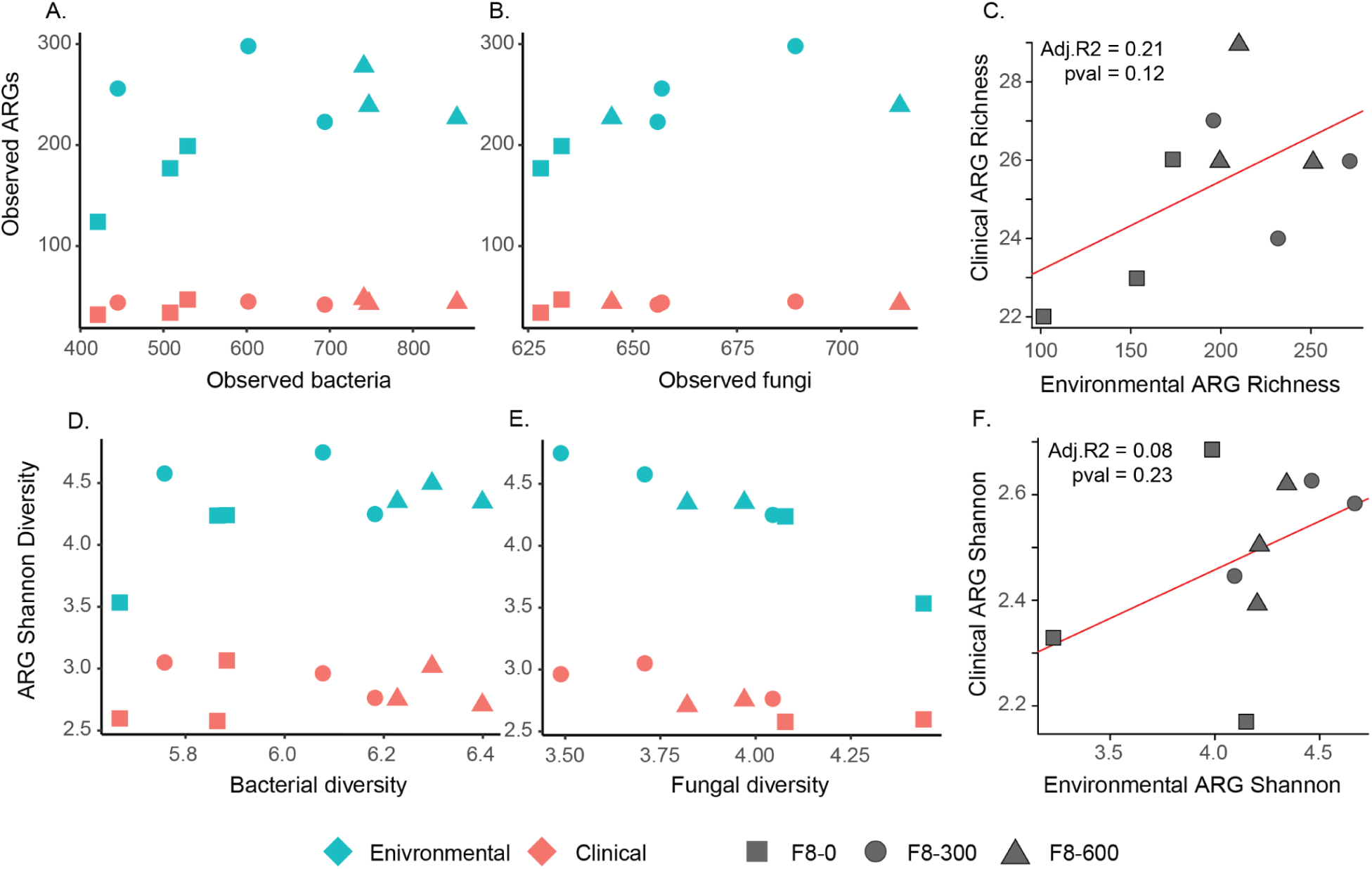
Correlations between antibiotic resistance genes diversity and microbial diversity. Total number of predicted environmental ARGs (blue) and clinical ARGs (red) are compared to **A**) the observed number of bacterial ASVs, **B**) the observed number of fungal ASVs, and **C**) the correlation between clinical and environmental ARG richness. Also, the Shannon diversity of predicted environmental ARGs (blue) and clinical ARGs (red) are compared to the diversity of **D**) the bacterial ASVs, **E**) fungal ASVs, and **F**) the correlation between clinical and environmental ARG diversity indexes.

However, unlike abundances, we did find that environmental ARG diversity positively correlated with bacterial diversity (*F*_(1,4)_ = 14.46; *P* = 0.01; Figure 4D) as well as with fungal diversity (*F*_(1,5)_ = 22.66; *P* = 0.005; Figure 4E). However, we did not find the same pattern when considering clinical ARG diversity relative to bacterial diversity (*F*_(1,4)_ = 0.12; *P* = 0.12; Figure 5D) and found that clinical ARG diversity slightly, but significantly, decreased as fungal diversity increased (*F*_(1,5)_ = 13.08; *P* = 0.01; Figure 4E). In other words, these results suggest that while there might be a positive relation between environmental ARG diversity and both bacterial and fungal diversity, this relation does not exist for clinical ARGs.

**Figure 5.**
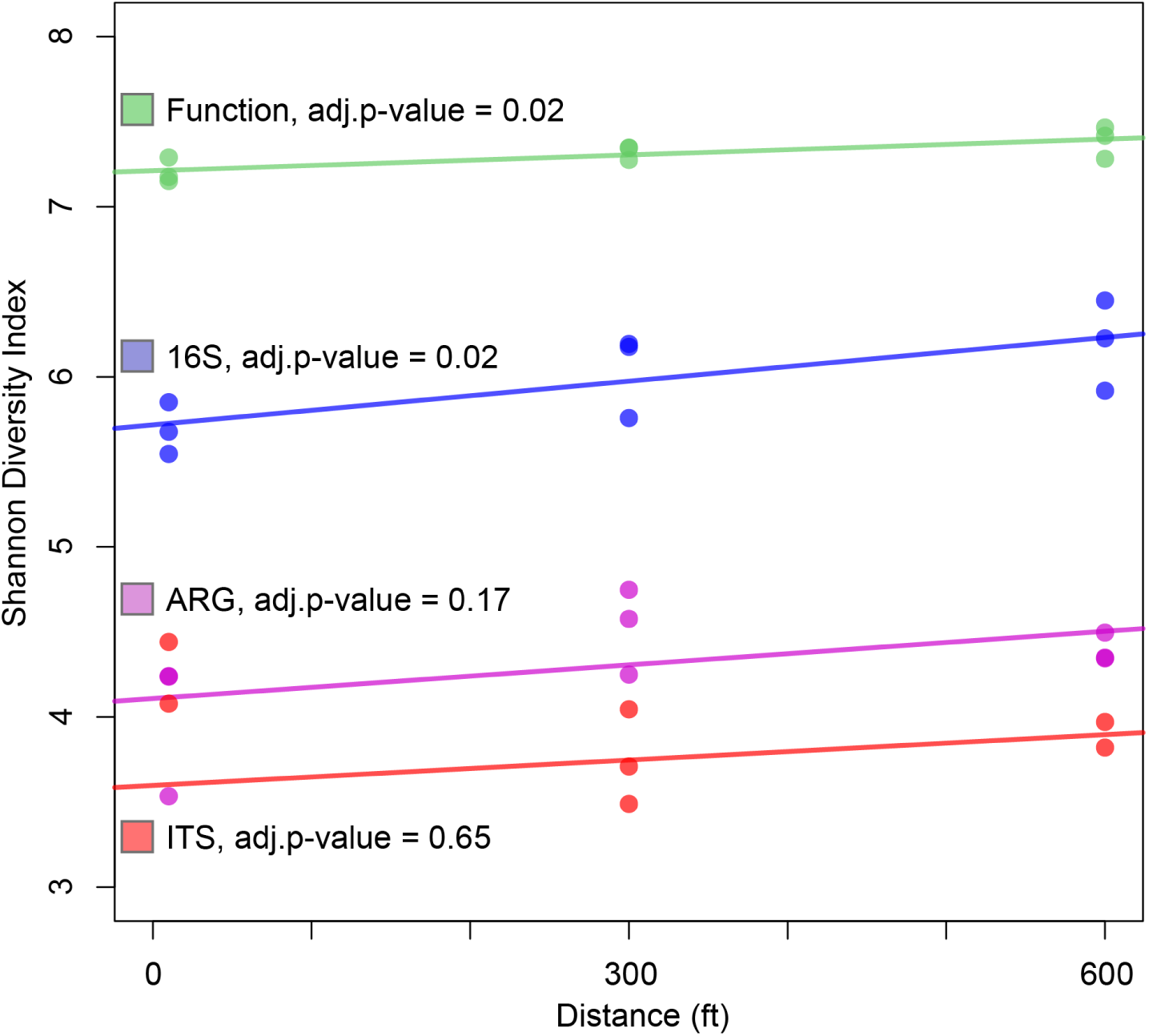
Linear trends of measured diversity indices over distance along the F8 transect. Linear trends, estimated using linear regression, are represented for diversity in bacterial ASVs (blue), fungal ASVs (red), predicted function genes (green), and antibiotic resistance genes (purple). Each point represents data from an individual sample.

Finally, we wanted to directly test the possibility that environmental ARGs had a direct correlation with clinical ARGs. However, we did find neither the Observed number of ARGs (*P* = 0.12, Figure 4C) nor the Shannon diversity indexes (*P* = 0.23, Figure 4F) are significantly correlated.

### 3.6 Correlations between ARGs with functional diversity

Since some genes may function in a way that encourages ARGs to arise and be selected for, we next sought to determine if any strong correlation may be observed between individual ARGs and gene functions. To do so, we performed a pairwise analysis of the correlation between relative gene abundances and relative ARG abundances. While some structure is present (Supplementary Figure S5), the vast majority of interactions (∼98.9%) were not statistically significant (*P*s > 0.05). Yet, we observed a few (∼1.1%) significant correlations (Benjamin-Hochberg adj-*P*s ≤ 0.05), including 4 clinically relevant ARGs (Supplementary Figure S5). The correlations included strongly positive relationships such as the cytochrome p450 involved in the cell stress response system with mexY, an important ARG in found pathogenic bacteria. We also found strongly positive correlations such as between the tetracycline resistance genes and the DNA-binding HTH domain TetR-type, a domain of proteins known for its role in the resistance to this antibiotic class. Taken together, our results suggest that the significant correlation between ARGs and gene diversity is not a spurious result of correlated abundance.

### 3.7 Investigating the effect of distance within the transect on microbial communities

When considering the general effect of the transect covering 600 feet of unused hay field, we observed a general trend suggesting that most indicators changed in relation to distance from the field’s edge. To test this hypothesis, we further investigated these trends using linear regression analyses. We found a significant positive linear relationship between distance from the creek and both functional and bacterial diversity (Figure 5). On the other hand, although slightly positive we did not find ARG or fungal diversity to have a significant linear relationship with distance.

## 4. Discussion

Bacterial infections resistant to antibiotics are an important public health issue and are predicted to become one of the main causes of mortality globally by 2050 (GBD 2021 Antimicrobial Resistance Collaborators 2024). Because antibiotic resistance genes (ARGs) are naturally present in microbial communities from environmental reservoirs (Perron, Whyte, et al. 2015; D’Costa et al. 2011, 2006)(Perron, Quessy, and Bell 2008; Despotovic et al. 2023)(Perron, Whyte, et al. 2015; D’Costa et al. 2011, 2006)(Perron, Quessy, and Bell 2008; Despotovic et al. 2023)(Perron, Whyte, et al. 2015; D’Costa et al. 2011, 2006), the World Health Organization as well as the Center for Disease Control and Prevention are advocating for a OneHealth approach that seeks to consider the important links between human health, animal health, and environmental health to manage the growing antibiotic-resistance crisis (McEwen and Collignon 2018). Thus, understanding the factors that shape the distribution of antibiotic resistance genes in the environment can provide key information to mitigate the emergence and spread of clinically relevant antibiotic resistance genes (Perron, Inglis, et al. 2015).

Here, we investigated how the diversity of antibiotic resistance genes related to bacterial and fungal diversity as well as total functional diversity in an old hay field along an agricultural transect located at the Hudson Valley Farm Hub, an active farm located in New York State. The aim of this study was to understand the role of bacterial and fungal diversity in shaping the presence of antibiotic resistance genes of different origin, environmental or clinical. To do so, we first established a baseline level by confirming our prediction that the total number of different genes observed in a sample correlated with overall bacterial and fungal diversity. In other words, the more bacterial and fungal types we found in a sample, the more functional genes we found on average. This enabled us to test whether trends in ARGs would follow the same general observation, i.e., increase with microbial and/or functional diversity, or would diverge from the latter.

When first looking at the total number of ARGs found in the environment, we found a positive correlation with bacterial, fungal, and functional diversity. This is not surprising as ARGs are indeed functions in themselves (Xu et al. 2024). In addition, many antibiotic resistance genes have additional functions in their natural environment such as molecule transport via the cell membrane (Walsh and Duffy 2013) or as signaling molecules at low concentrations (Yim, Wang, and Davies 2007). Interestingly, when investigating if any specific functional category of gene could explain the distribution of ARGs, we found that the vast majority of functions (>98%) were not significantly correlated with ARGs, suggesting that overall functional diversity rather than a focus on a specific group of genes (or possibly taxa) is the best indicator for the presence of high ARG diversity in the environment. These results confirm not only that total ARGs can be extensive in the environment, but also that they could even be considered a measure of soil health. While unconventional, this could be useful given the increasing availability of pipelines to document the presence of antibiotic resistance in metagenomics data (McArthur et al. 2013; Hackenberger et al. 2024; “[No Title],” n.d.; Olatunji et al. 2024).

However, when focusing on ARGs of clinical importance, we found that the relationship between the latter and functional diversity was only marginally significant at best when considering the total number of genes, and disappeared altogether when considering the relative abundance of each gene. In other words, we did not observe an increase in clinically-relevant ARGs solely because we observed an increase in gene functions, suggesting that other mechanisms exist that shape the distribution of such genes in soil environments. Indeed, we found that ARGs of clinical importance were not correlated with an increase in functional diversity nor with bacterial diversity. This result is interesting given that bacteria are the predominant carrier of antibiotic resistance genes. At least two mechanisms could explain such a finding. For one, many clinical resistance genes spread into clinical pathogens after being mobilized on mobile genetic elements such as plasmids, phages, and integrons thus spreading horizontally under selective pressures imposed by higher concentration (Larsson and Flach 2022; Barathe et al. 2024). Importantly, such results apply to agricultural soils (Osbiston, Oxbrough, and Fernández-Martínez 2021). Second, many antibiotic resistance genes also found their way into clinical settings by being acquired by opportunistic pathogens that tend to show higher adaptive abilities like exogenous DNA acquisition (Liang et al. 2020; Sanz-García et al. 2021; Chng et al. 2020). While the two mechanisms are not mutually exclusive, both make it possible that changes in ARG distribution in response to local selective pressures could be significantly different than that of their bacterial hosts.

In addition, we also found that ARG diversity was negatively correlated with fungal diversity (Figure 5E). The widespread capacity of fungi to produce natural antibiotics has been continuously explored (Zhao et al. 2019)(Silber et al. 2016) since the discovery of penicillin (Fleming, 1929). For that reason, it is often predicted that bacteria exposed to a diversity of antibiotic-producing fungi would harbor a higher number of antibiotic-resistance genes (Nazir et al. 2017)(X. Wang et al. 2025). Moreover, fungi can often carry ARGs themselves and are known to actively help the horizontal transfer of ARGs between bacteria (Nazir et al. 2017)(M. Zhang et al. 2014), further increasing the expected level of ARG diversity in the presence of fungi. However, our findings challenge this idea. Notably, this surprising result is consistent with recent research that finds that fungi can change resistome structure (Y.-F. Wang et al. 2021), and that bacteria and fungi can present widespread antagonistic behavior in soil depending on the local environment (Bahram et al. 2018). It is also possible that in the sites with higher diversity of fungi, competition may be selecting bacteria with specific resistance genes rather than a generally diverse resistome (Toprak et al. 2011). Given the limitation of *ITS* amplicon sequencing to document fungal taxonomy, future work, combining a culture-based approach and genomics, will likely be needed to confirm this hypothesis.

While our results suggest that promoting diverse fungal communities could be used to reduce the distribution of ARGs in the environment, our current study does not allow us to identify under what conditions this relationship holds. Fungal communities are known to be influenced by a complex interplay of factors such as soil properties, environmental conditions, and biotic interactions (Nazir et al. 2017). For example, fungal communities can be more sensitive than bacterial communities to environmental variables like precipitation or nutrient availability (He et al. 2023). Furthermore, many fungi can have symbiotic relationships with plants, meaning that their distribution will be codependent (Lanfranco, Bonfante, and Genre 2016). Given that plant diversity changes over different sections of the hay field, it is likely that fungal diversity was affected by other environmental factors at these sites. We plan to investigate the interaction between plant diversity, fungal diversity, and antibiotic-resistance genes in future work.

It is also possible that we did not find correlations between specific fungal taxa and ARGs due to the fact that we used a single-gene amplicon library strategy to estimate diversity. While this technique for studying microbiomes has been useful in a diverse array of studies, it is known that amplicon library sequencing underestimates genetic variability at the sub-genus level (Straub et al. 2020); (Thines et al. 2018). The latter may be especially relevant with antibiotic resistance genes, which are often exchanged horizontally within and among species (Guernier-Cambert et al. 2021). The fact that antibiotic resistance gene diversity correlated with functional diversity, which used WGS, indicates that novel approaches, such as long-read sequencing, to estimate bacteria diversity might be better suited for such work in the future.

Finally, while the transect was originally designed to capture different ecological niches in relation to the field edges, our results suggest that there exists a disturbance gradient along the transect. Using heavy metal concentrations, we found that, on average, site F8-0 presented the highest concentration of metals, which then decreased as the transect moved into the fallow. The additional site considered in our study, F8-W, located between the road and the Esopus Creek also presented high levels of heavy metals, similar to site F8-0, with the exception of copper levels being slightly lower and lead level being higher than the site near the field’s edge. While both metals are associated with vehicle movement, copper is known to be more reactive in the environment and could indicate that the site near the water is more active. Interestingly, we also found that site F8-W also showed clear signs of disturbance with an increased level of Proteobacteria compared to the other sites (Watkinson et al. 2009). Similarly, this site also presents a high diversity of clinically-relevant ARGs, which is not surprising given that the Esopus Creek has been shown to contain harmful levels of pathogenic bacteria from time to time (“Hudson River Estuary Data” 2010). Because some ARGs can confer cross-resistance to heavy metals (Gillieatt and Coleman 2024), this could explain, at least in part, the fact that clinical ARGs were higher near the road. Indeed, *aadA*, *bla*CTX-M, and *vanA* have all been shown to be co-selected with the presence of heavy metals, especially copper in agricultural settings (Seiler and Berendonk 2012; Wales and Davies 2015). Therefore, it is possible that the presence of the Esopus Creek and the apparent gradient in heavy metals concentrations could provide a confounding explanation for the selection of antibiotic resistance correlating with microbial biodiversity. Future work will be necessary to determine what mechanisms are more important.

Taken together, our results suggest that land use, specifically agricultural practices, that impact fungal diversity can result in changes in the diversity of clinically relevant ARGs. Notably, this correlation is not observed between clinically relevant ARGs and bacterial diversity, suggesting that diversity of clinically relevant ARGs is not solely due to the bacterial population, as is the case with environmental ARGs. This suggests that farming practices that favor fungal diversity could result in the reduction of antibiotic resistance genes spread in the environment. Whether such results could be applied to other environments such as natural habitats that serve as buffers remains to be investigated.

## 5. Data accessibility

All computational scripts used in this analysis of the data are publicly available as a versioned GitHub repository here: https://github.com/pspealman/transect/releases/tag/v0.1

## Acknowledgments

This study is part of the Applied Farmscape Ecology Research Collaborative (AFERC). AFERC is co-coordinated by Hawthorne Valley Farmscape Ecology Program and The Hudson Valley Farm Hub and is funded by The Hudson Valley Farm Hub. The study was also funded by the Bard Summer Research Institute of Bard College. The authors would like to thank Christopher Benincasa, Hannah Herrick, and Tejaswee Neupane for their assistance in the field and DNA extraction and Chad McKinney for his work on qPCR.

## Supplemental Materials

**Figure S1.**
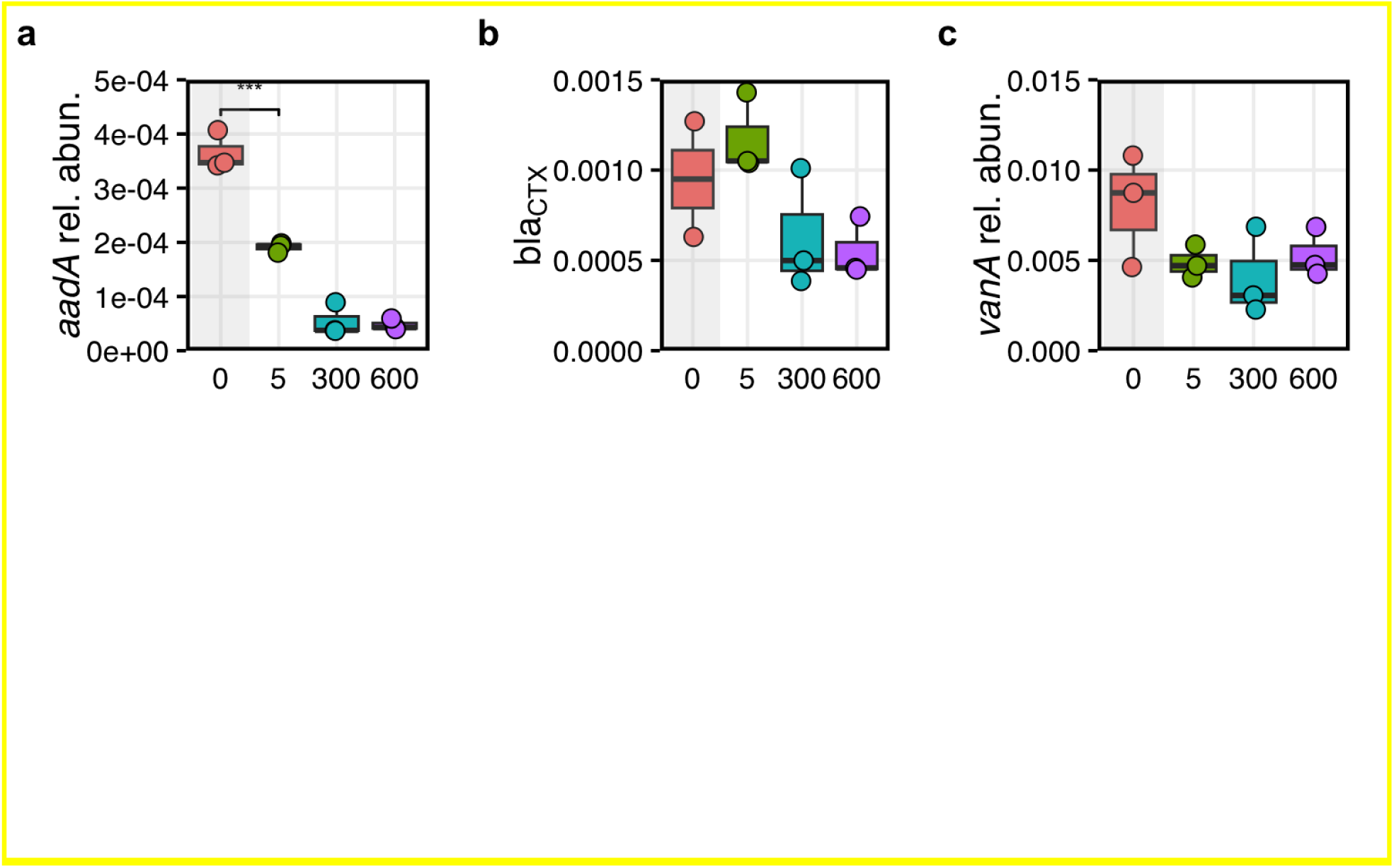
Relative abundance of three antibiotic resistance genes along the F8 agricultural transect. Relative abundance for antibiotic resistance genes **a**) *aadA*, **b**) *blaCTX*, and **c**) *vanA* was estimated by dividing the gene copy number by the copy number of *16S rRNA* gene identified in each sample using quantitative PCR. We used ANOVA to compare r.a. between site F8-W and the transect sites while we used linear regression analyses to assess the trend observed in the transect, i.e., excluding F8-W. We found a significant decrease in relative abundance for *aadA* and *blaCTX* and no statistically significant trend for *vanA*. Each dot represents an individual replicate collected at each site and whiskers represent quartiles.

**Figure S2.**
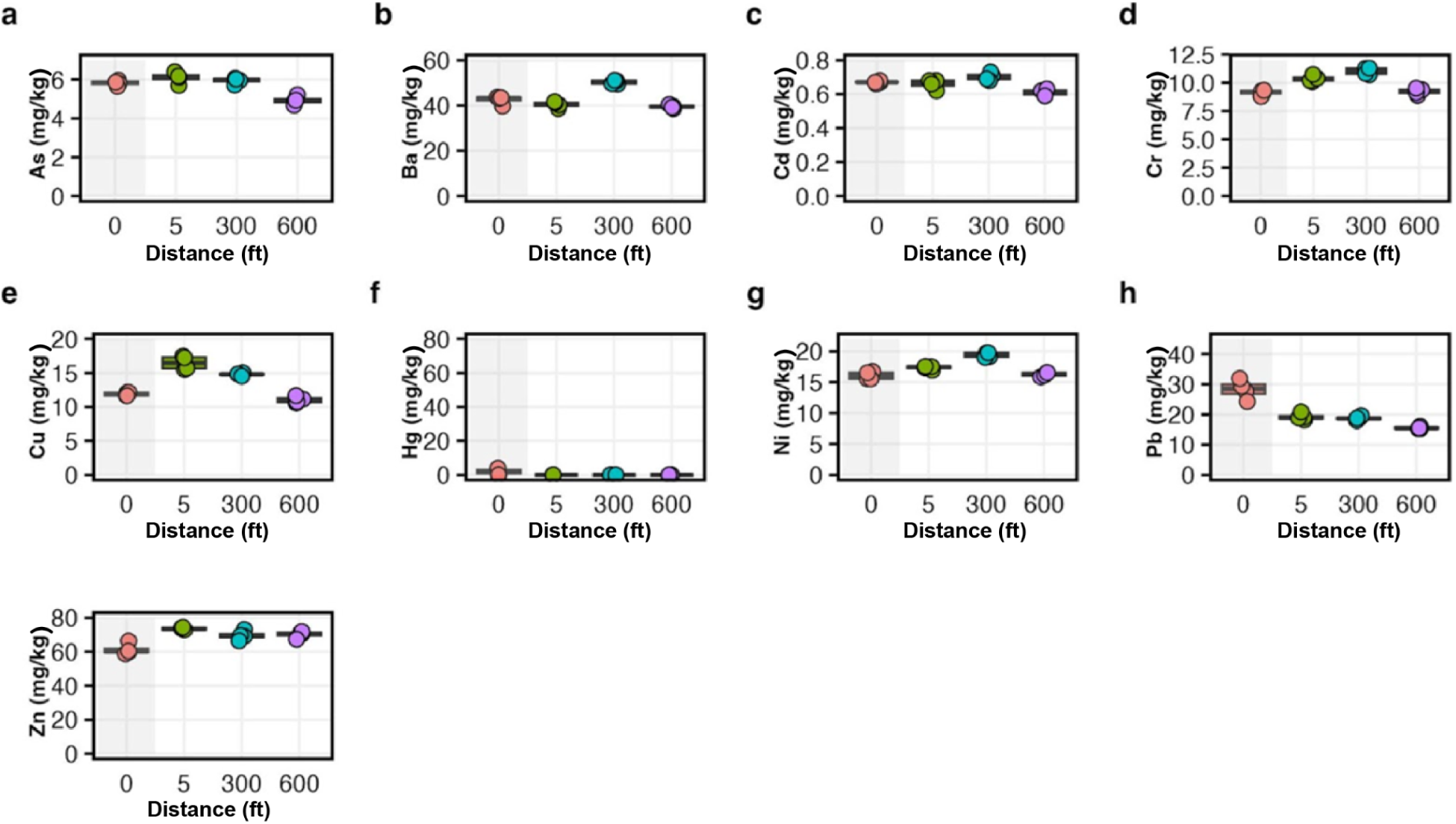
Concentration of nine heavy metals along the F8 agricultural transect. Concentrations (mg/kg) trends over distance (ft) were estimated using two methods for **a**) Arsenic (As), **b**) Barium (Ba), **c**) Cadmium (Ca), **d**) Chromium (Cr), **e**) Copper (Cu), **f**) Mercury (Hg) **g**) Nickel (Ni), **h**) Lead (Pb), and **i**) Zinc (Zn). We used ANOVA to compare concentrations between site F8-W and the transect sites while we used linear regression analyses to assess the trend observed in the transect, i.e., excluding F8-W. Each dot represents an individual replicate collected at each site.

**Figure S3.**
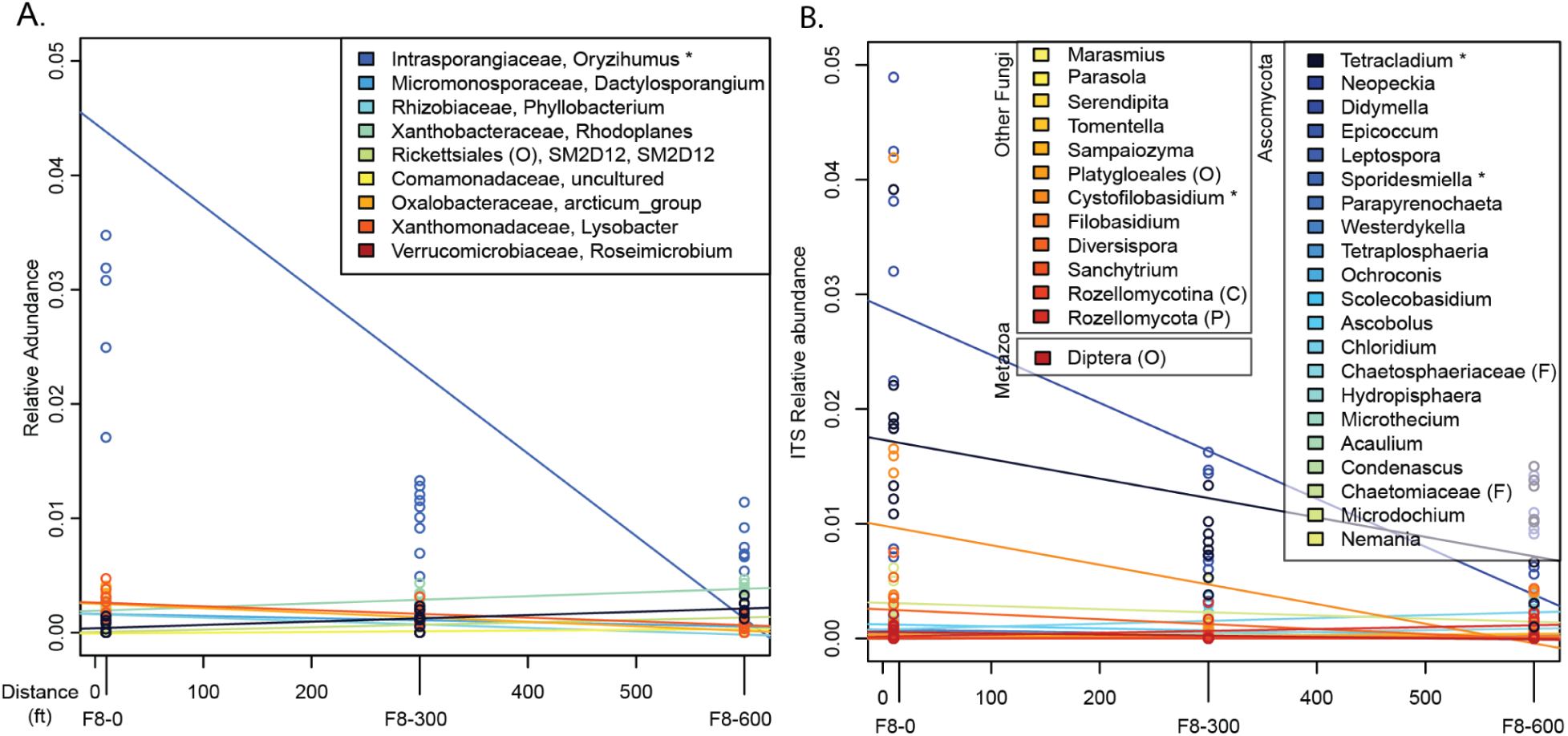
Abundance of genera as a function of distance. (A) We used a linear model of the relative abundance of bacterial genera as a function of distance and only found one genera, *Oryzihumus,* with a relative abundance greater than 1% that significantly changed as a function of distance. (B) Similarly for fungal genera, we found 3 genera with relative abundances greater than 1% significantly changed over distance from the water; the Ascomycota *Sporidesmiella* and *Tetracladium*, and the Basidiomycota *Cystofilobasidium* (B).

**Figure S4.**
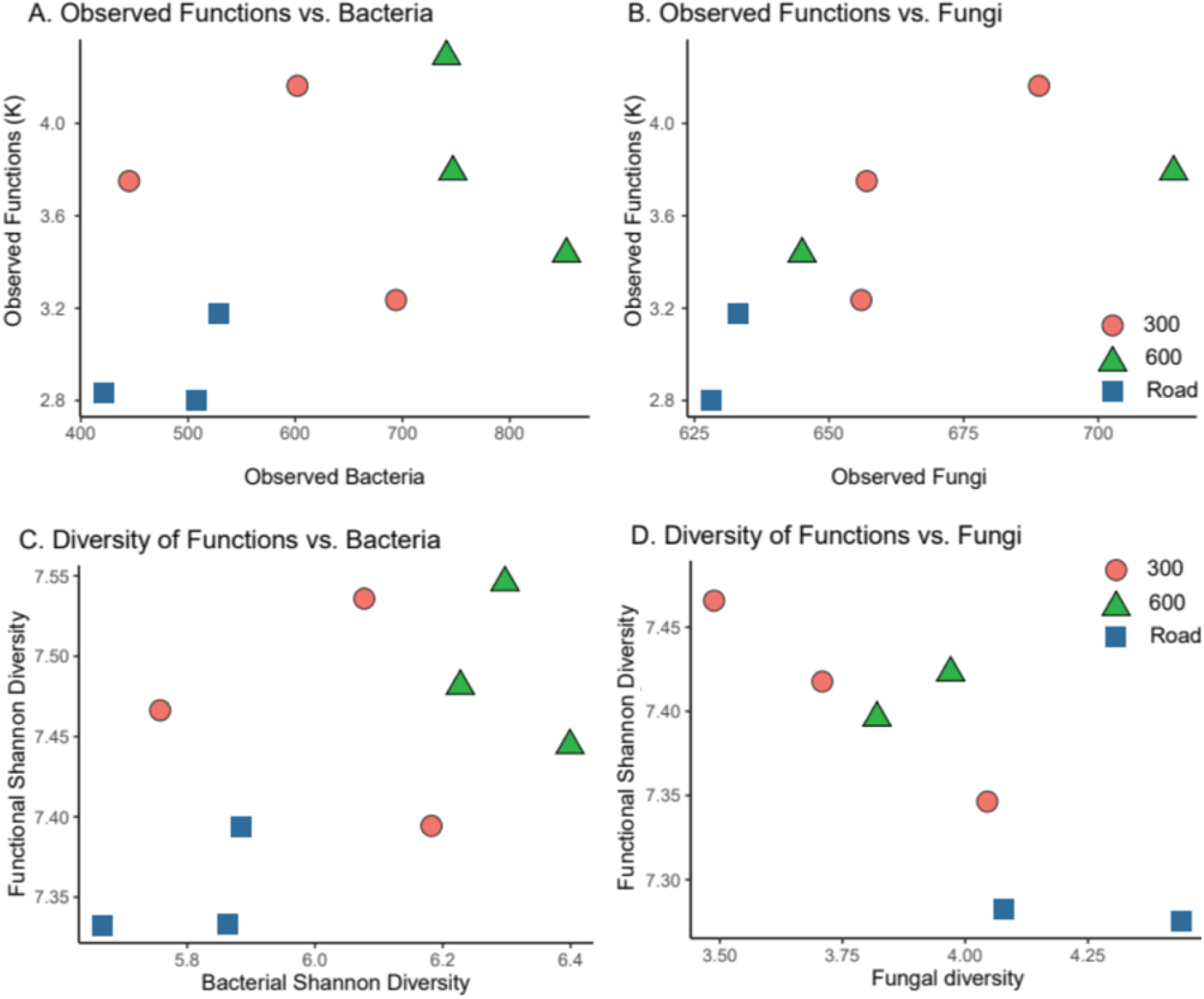
Correlation between diversity of functions identified and microbial communities. (**A**) We find that the total number of observed predicted functions did not significantly increase with the number of observed bacterial ASVs (*F*_(1,6)_ = 3.23; *R*^2^ = 0.56, *P* = 0.12) (**C**) nor that the diversity (H) of functional genes correlated with bacterial diversity (*F*_(1,6)_ = 0.09; *R*^2^ = 0.46, *P* = 0.77). (**B**) we found that the total number of functions did not significantly correlate with the number of observed fungal ASVs (*F*_(1,4)_ = 3.52; *R*^2^ = 0.45, *P* = 0.13), but that (**D**) functional diversity measured as Shannon Diversity Index, was negatively correlated with fungal diversity (*F*_(2,2)_ = 316.12; *R*^2^ = 0.99, *P* = 0.003) suggesting that fungal diversity could play an important role in shaping functional diversity.

**Figure S5.**
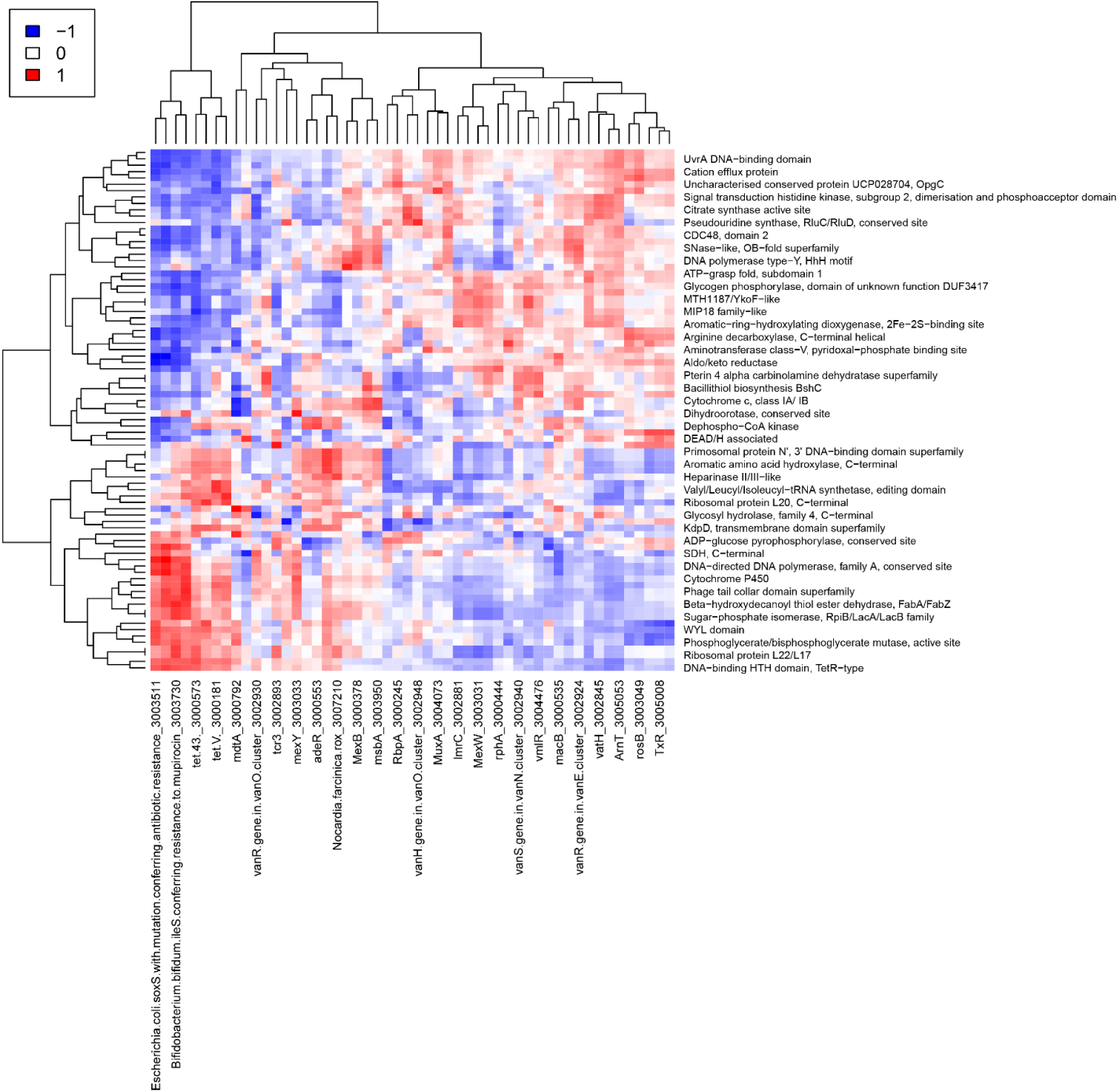
Correlation between antibiotic resistance genes and predicted gene functions. Heatmap shows the significantly correlated (R-value) of the relative abundances of functions and ARGs estimated using linear regression, i.e., Benjamin-Hochberg adj-*P*s <= 0.01. Each function and ARG must be observed in a minimum of six samples to be included. Four clinical ARGs are present, macB, MexB, MexW, MexY.

### Supplemental Tables

**Table S1.** Tab delimited file. Contains sequencing run quality control numbers for amplicon sequencing runs generated by denoising using DADA2.

**Table S2.** Tab delimited file. Contains QIIME2 Feature IDs with taxonomic assignments and confidence scores.

**Table S3.** Tab delimited file. Contains omnibus and pairwise alpha-diversity tests (Evenness, Faith-PD, Shannon) and beta-diversity tests (Jaccard, Bray-Curtis, Unweighted UniFrac, Weighted UniFrac) for both ITS (rarefaction = 350K) and 16S (rarefaction = 10K) runs.

**Table S4.** Tab delimited file. Contains all pairwise Bray-Curtis dissimilarity (ie. percentage difference) for all 16S and ITS samples.

**Table S5.** Condition for antibiotic resistance gene qPCR.

